# *Listeria monocytogenes* Requires Phosphotransferase Systems to Facilitate Intracellular Growth and Virulence

**DOI:** 10.1101/2024.08.12.607557

**Authors:** Matthew J. Freeman, John-Demian Sauer

## Abstract

The metabolism of bacterial pathogens is exquisitely evolved to support growth and survival in the nutrient-limiting host. Many bacterial pathogens utilize bipartite metabolism to support intracellular growth by splitting carbon utilization between two carbon sources and dividing flux to distinct metabolic needs. For example, previous studies suggest that the professional cytosolic pathogen *Listeria monocytogenes* (*L. monocytogenes*) utilizes glycerol and hexose phosphates (e.g. Glucose-6-Phosphate) as catabolic and anabolic carbon sources in the host cytosol, respectively. However, the role of this putative bipartite glycerol and hexose phosphate metabolism in *L. monocytogenes* virulence has not been fully assessed. Here, we demonstrate that when *L. monocytogenes* is unable to consume either glycerol (Δ*glpD*/Δ*golD*), hexose phosphates (Δ*uhpT*), or both (Δ*glpD*/Δ*golD*/Δ*uhpT*), it is still able to grow in the host cytosol and is minimally attenuated *in vivo* suggesting that *L. monocytogenes consumes alternative carbon source(s) in the host. An in vitro* metabolic screen using BioLog’s phenotypic microarrays demonstrated that both WT and PrfA* *L. monocytogenes, a strain with constitutive virulence gene expression mimicking cytosolic replication, use phosphotransferase system (PTS) mediated carbon sources. These findings contrast with the existing metabolic model that cytosolic L. monocytogenes* expressing PrfA does not use PTS mediated carbon sources. We next demonstrate that two independent and universal phosphocarrier proteins (PtsI [EI] and PtsH [HPr]), essential for the function of all PTS, are critical for intracellular growth and virulence in vivo. Finally, virulence phenotypes of these mutants were additive to mutants unable to consume glycerol and hexose phosphates (Δ*glpD*/Δ*golD*/Δ*uhpT*) *in vivo*, suggesting that hexose phosphates and glycerol are relevant metabolites *in vivo* in addition to those derived from PTS. Taken together, these studies indicate that PTS are critical virulence factors for the cytosolic growth and virulence of *L. monocytogenes*.

## INTRODUCTION

The mammalian cytosol is a stringent and hostile environment that restricts the growth of bacteria not specifically adapted to that niche (1–5). One mediator of bacterial growth restriction in the host cytosol is nutrient availability whereby cells actively or passively limit access to vital nutrient resources preventing bacterial growth. For example, intracellular pathogens are metabolically restricted through limited access to metal ions, vitamin cofactors, and amino acid pools (6–9). More specifically, macrophages contribute to this metabolic foray by shifting metabolism between M1 or M2 states, altering concentrations of metals, and rewiring glycolysis and the tricarboxylic acid cycle to control bacterial pathogens (10–12). Despite this well-orchestrated defense, canonical cytosolic pathogens such as *Listeria monocytogenes* (*L. monocytogenes*) can replicate in this environment at a rate equivalent to that in rich media (13). This rapid growth allows *L. monocytogenes* to disseminate to distant sites of infection from the intestine (spleen, liver, & meninges), overtake host defenses thereby delaying antibiotic treatment, and culminating in a mortality rate approaching 30% (14–16). Defining bacterial metabolism can reveal novel targets for antibiotics and a better understanding of host-pathogen interactions. Despite significant progress, there are significant unknowns about what metabolites pathogens such as *L. monocytogenes* are using in their respective host environments, how these nutrients are acquired, and what impact this has on the host response to infection (17–20). Key challenges to progress in understanding host-pathogen metabolisms include an incomplete understanding of the metabolic profile of host cells during infection and redundant metabolic pathways in both the host and the pathogen.

In addition to *L. monocytogenes’* role as an important pathogen, it serves as a powerful model organism to answer questions about host-pathogen interactions (21). *L. monocytogenes*’ virulence genes and intracellular lifecycle have been well studied including in macrophages which serve as a primary replicative niche and host dissemination vehicle (13,22). *L. monocytogenes* encodes the master transcriptional virulence regulator, PrfA, that controls many key virulence factors to mediate the *L. monocytogenes* cytosolic lifecycle including Listeriolysin O (LLO) that facilitates cytosolic access, the phospholipase Cs (PLCs) that contribute to secondary vacuole escape, ActA which mediates intracellular motility and cell-to-cell spread, and the hexose phosphate transporter UhpT which contributes to cytosolic metabolism (13,22). PrfA is activated upon entry into the cytosol via allosteric regulation by glutathione, however PrfA* mutations such as the G145S mutation result in constitutive virulence gene expression and upregulation of *uhpT* thereby mimicking *L. monocytogenes’* cytosolic lifestyle (23,24).

Finally, well-defined *ex vivo* and *in vivo* models of *L. monocytogenes* infection and its genetic tractability allow for a thorough examination of how metabolic perturbation might impact its virulence and host response to that metabolic perturbation (25).

Previously, *L. monocytogenes* has been employed as a tool to understand bacterial cytosolic metabolism through isotopologue analysis. Specifically, this work examined the metabolism of *L. monocytogenes* during the infection of macrophages, a necessary step in disseminated infection *in vivo* (26,27). The Eisenreich and Goebel groups previously identified glycerol and hexose phosphates (e.g. Glucose-6-Phosphate) as the primary carbon metabolites used by *L. monocytogenes* during cytosolic replication (28–31). These findings were supported by the fact that *Listeria innocua*, a non-pathogenic *Listeria* species, lacks the transporter necessary for use of hexose phosphates (UhpT) and that glycerol is a common metabolite used by a variety of intracellular pathogens (32,33). Further, these analyses suggested that glycerol and hexose phosphates are primarily funneled into lower glycolytic catabolism and pentose phosphate pathway anabolism, respectively. Despite this, *L. monocytogenes* mutants lacking the ability to use glycerol (Δ*glpD*) or hexose phosphates (Δ*uhpT*) individually maintain intracellular growth and significant virulence (28,31). The lack of robust virulence phenotypes could be due to incomplete perturbation of specific carbon source use, a lack of physiologic relevancy of the *ex vivo* models, or the ability of *L. monocytogenes* to utilize alternative, yet to be defined carbon sources (34–38).

In addition to being able to replicate in the host cytosol following infection, *L. monocytogenes* also lives as a saprophyte in the environment and in food production facilities where its metabolic potential has also been studied intensely (39–41). Like many other bacteria, *L. monocytogenes* can utilize phosphotransferase systems (PTS) in these environments to acquire free sugar (42). The *L. monocytogenes* 10403s strain used in this study encodes 29 complete PTS, encoded by a collection of 86 genes (38,43,44). Interestingly, other *L. monocytogenes* strains as well as different *Listeria* species such as *L. innocua* and *L. welshimeri* vary in the specific PTS they encode suggesting flexibility in the use of PTS (45–48). Early work has shown these differences may be important for virulence, but a global understanding of their importance remains unknown (45,48,49). Mechanisms of PTS function are well-defined and reviewed elsewhere (50); yet, there are many open questions about the relationship between PTS components’ structure, sugar specificity, regulatory inputs/outputs, and ultimately their importance during infection (50). PTS mediate carbon source import and phosphorylation, with secondary functions on transcriptional regulation having been reported (51). PTS import free sugars following binding of a sugar to pre-phosphorylated import permeases. This pre-phosphorylation is tied to the lower glycolytic conversion of phosphoenolpyruvate (PEP) to pyruvate through two phosphocarrier proteins (PtsI [EI] & PtsH [HPr-His]). The result of this phospho-cycling is that free sugars are phosphorylated during import and readied for direct funneling to glycolysis. It has previously been reported that PrfA-dependent virulence gene expression is repressed when *L. monocytogenes* is utilizing primarily PTS-dependent carbon sources (41,52,53). Conversely, when PTS function is blocked via deletion of the HPr phosphocarrier protein (Δ*ptsH*), PrfA-dependent virulence gene expression is significantly increased *in vitro* (20,31,53,54). Because of these observations, PTS are thought to be inactive during *L. monocytogenes* intracellular growth and virulence (43). Instead, the reported PTS-mediated *prfA* downregulation has led to a widespread belief that PTS are inactive during infection (38,44,55,56).

In this work, we assessed the contribution of glycerol and hexose phosphate metabolism to *L. monocytogenes* virulence *in vivo* finding that glycerol and hexose phosphates are neither sufficient nor essential to support the intra-macrophage growth and only minimally contribute to virulence of *L. monocytogenes*. A metabolic screen of carbon sources revealed highly similar metabolite utilization between WT and PrfA* *L. monocytogenes,* including the use of PTS mediated carbon sources. Ablation of all PTS-mediated carbon acquisition via deletion of the phosphocarrier proteins EI (Δ*ptsI*) or HPr (Δ*ptsH*) revealed that *L. monocytogenes* requires PTS function to support intracellular growth and virulence. Finally, the phenotypes associated with loss of PTS function are additive with the inability to utilize glycerol and hexose phosphates (Δ*glpD*/Δ*golD/*Δ*uhpT/ΔptsI or* Δ*glpD*/Δ*golD/*Δ*uhpT/ΔptsH*) demonstrating that *L. monocytogenes* uses a highly complex and varied network of metabolites to promote rapid intracellular replication during infection.

## RESULTS

### *glpD*/*golD* and *uhpT* genes are required for *L. monocytogenes* to consume glycerol and hexose phosphate, respectively

*L. monocytogenes* uses host-derived glycerol and hexose phosphates during cytosolic replication, as defined using isotopologue metabolomics analysis by the Goebel and Eisenreich labs (28,30,31), however whether these are the predominant carbon sources used during *in vivo* infection has not been fully explored. Although *uhpT, glpD* and *golD* mutants have been studied in isolation, the phenotype of combination metabolic mutants of parallel glycerol utilization pathways (Δ*glpD*/Δ*golD*) and of glycerol with hexose phosphate pathways (Δ*glpD*/Δ*golD/*Δ*uhpT*) have not been assessed. We aimed to test the hypothesis that combination metabolic mutants of Δ*glpD*/Δ*golD*, Δ*uhpT*, and combination Δ*glpD*/Δ*golD/*Δ*uhpT* would completely ablate *L. monocytogenes* growth on glycerol, hexose phosphates, and both, respectively, To do this we generated these metabolic mutants and assessed growth in *Listeria* synthetic media (LSM) with glucose, glycerol, glucoe-6-phosphate (+glutathione), or glycerol/glucose-6-phosphate (+glutathione) as the sole carbon source (**Fig. 1A** and **S1 Fig.**). LSM with defined sole carbon sources was inoculated with WT *L. monocytogenes* or the indicated mutants, and growth was assessed via OD_600_ absorbance every 15 minutes. Importantly, any LSM containing hexose phosphates required further supplementation with 10 mM reduced glutathione to induce *prfA,* and therefore *uhpT*, expression (23,57). Of note, we found that a Δ*glpD* mutant was still able to grow in LSM supplemented with glycerol and a second glycerol utilization pathway required ablation (Δ*golD*) to limit *in vitro* growth (data not shown) (37). We found that glycerol (Δ*glpD*/Δ*golD*), hexose phosphate (Δ*uhpT*), and combined (Δ*glpD*/Δ*golD/*Δ*uhpT*) mutants were not defective for *in vitro* growth when compared to WT *L. monocytogenes* in LSM with glucose as the sole carbon source (**Fig. 1B and S1A Fig.**). In contrast, when Δ*glpD*/Δ*golD* or Δ*uhpT* mutants were supplied with glycerol and hexose phosphates, respectively, they were unable to grow while WT *L. monocytogenes* sustained growth (**Fig 1C**). Mutants lacking only the hexose phosphate transporter (Δ*uhpT*) showed sustained growth in LSM with glycerol alone, but they were only able to growth modestly in LSM with glycerol and hexose phosphates (**S1B-1D Fig.**). Finally, a mutant defective for glycerol and hexose phosphate (Δ*glpD*/Δ*golD/*Δ*uhpT*) utilization was fully capable of growing in defined media with glucose as the carbon source but was unable to grow on glycerol, hexose phosphates, or glycerol/hexose phosphates combined when compared to WT *L. monocytogenes* (**Fig 1B and 1C**). Taken together, these data demonstrate that there are not additional unknown glycerol or hexose-phosphate utilization pathways and that the Δ*glpD*/Δ*golD/*Δ*uhpT* mutant is incapable of growing on glycerol and hexose phosphates as primary carbon sources. Similarly, the Δ*glpD*/Δ*golD/*Δ*uhpT* mutant shows no growth defects when supplied with glucose as its sole carbon source.

**Figure 1.**
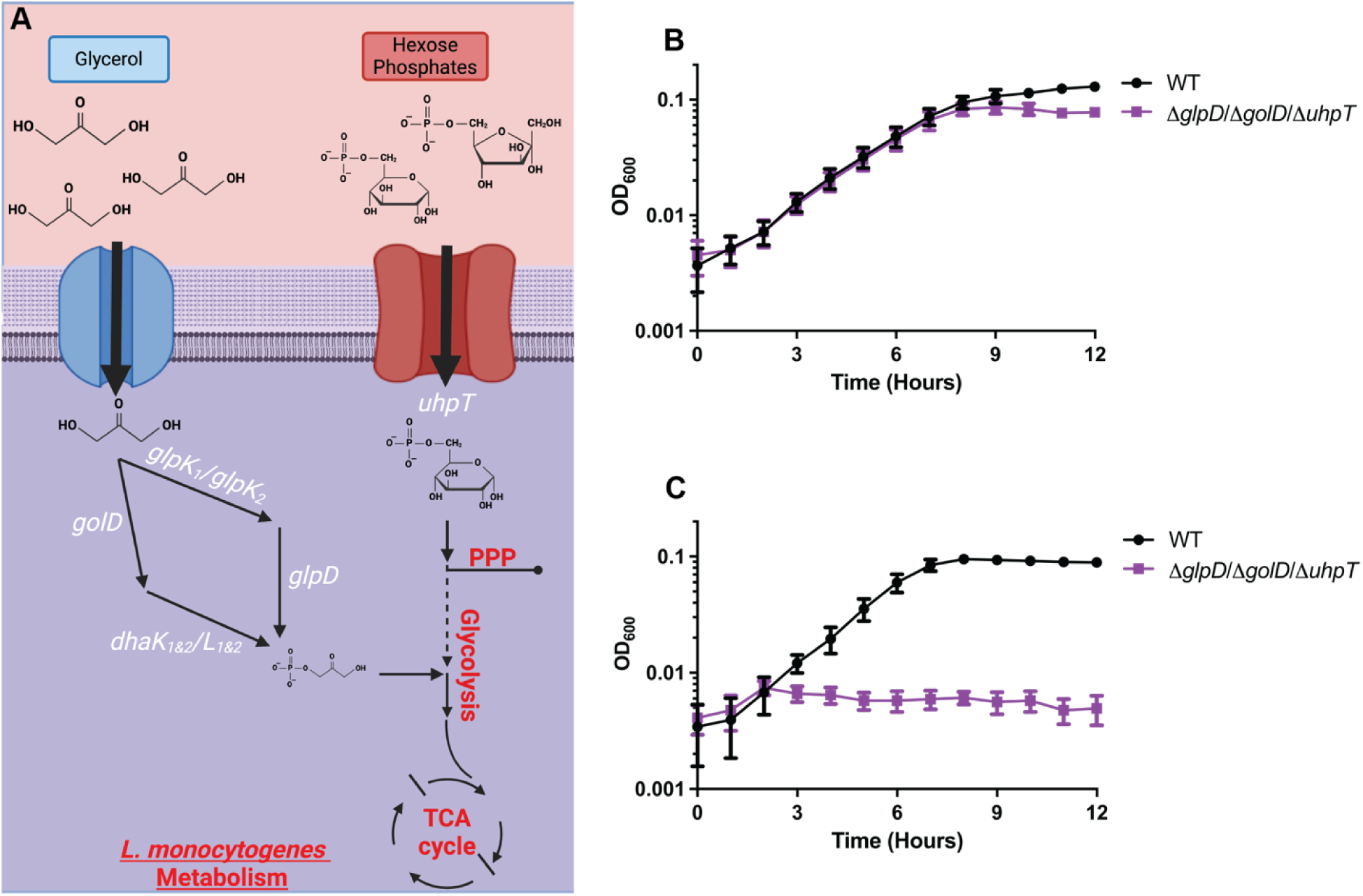
Δ*glpD*, Δ*golD* and *uhpT* are required for growth on glycerol and hexose phosphates. (A) Simplified model of glycerol being imported and funneled into two parallel glycerol utilization pathways (*glpD* and *golD*) for entry into central catabolic glycolysis. Hexose phosphate imported and funneled into the anabolic pentose phosphate pathway. Indicated strains were grown in LSM at 37°C, shaking at 250 r.p.m. with the addition of 55mM glucose (B) or carbon equivalent amounts of hexose phosphates (+10mM glutathione) and glycerol (C). OD_600_ was monitored every 15 minutes for 12 hours. Data represents average of three technical replicates from one representative of three biological replicates.

### Δ*glpD*/Δ*golD/*Δ*uhpT L. monocytogenes* mutants unable to use hexose phosphates and glycerol replicate in macrophages and are mildly attenuated *in vivo*

Glycerol and hexose phosphates have been demonstrated via isotopologue metabolomic flux analysis to be utilized by *L. monocytogenes* during intra-macrophage replication (29,30). We hypothesized that mutants deficient in both pathways would be attenuated for virulence. To determine if glycerol, hexose phosphates, or both were necessary for cytosolic replication of *L. monocytogenes* we performed intra-macrophage growth curves in murine bone marrow-derived macrophages (BMDMs). Intracellular growth curves in BMDM demonstrated that individual (Δ*glpD*/Δ*golD* and Δ*uhpT*) and combined (Δ*glpD*/Δ*golD/*Δ*uhpT*) metabolic mutants unable to use glycerol and/or hexose phosphates were still able to grow in the host cytosol with kinetics like WT *L. monocytogenes* (**Fig 2A)**. This data suggests that in a single cycle infection in primary BMDMs, *L. monocytogenes* does not require glycerol or hexose phosphate to support cytosolic replication. Further, this data suggests that *L. monocytogenes* must be able to use alternate undefined carbon source(s) to support cytosolic growth.

**Figure 2.**
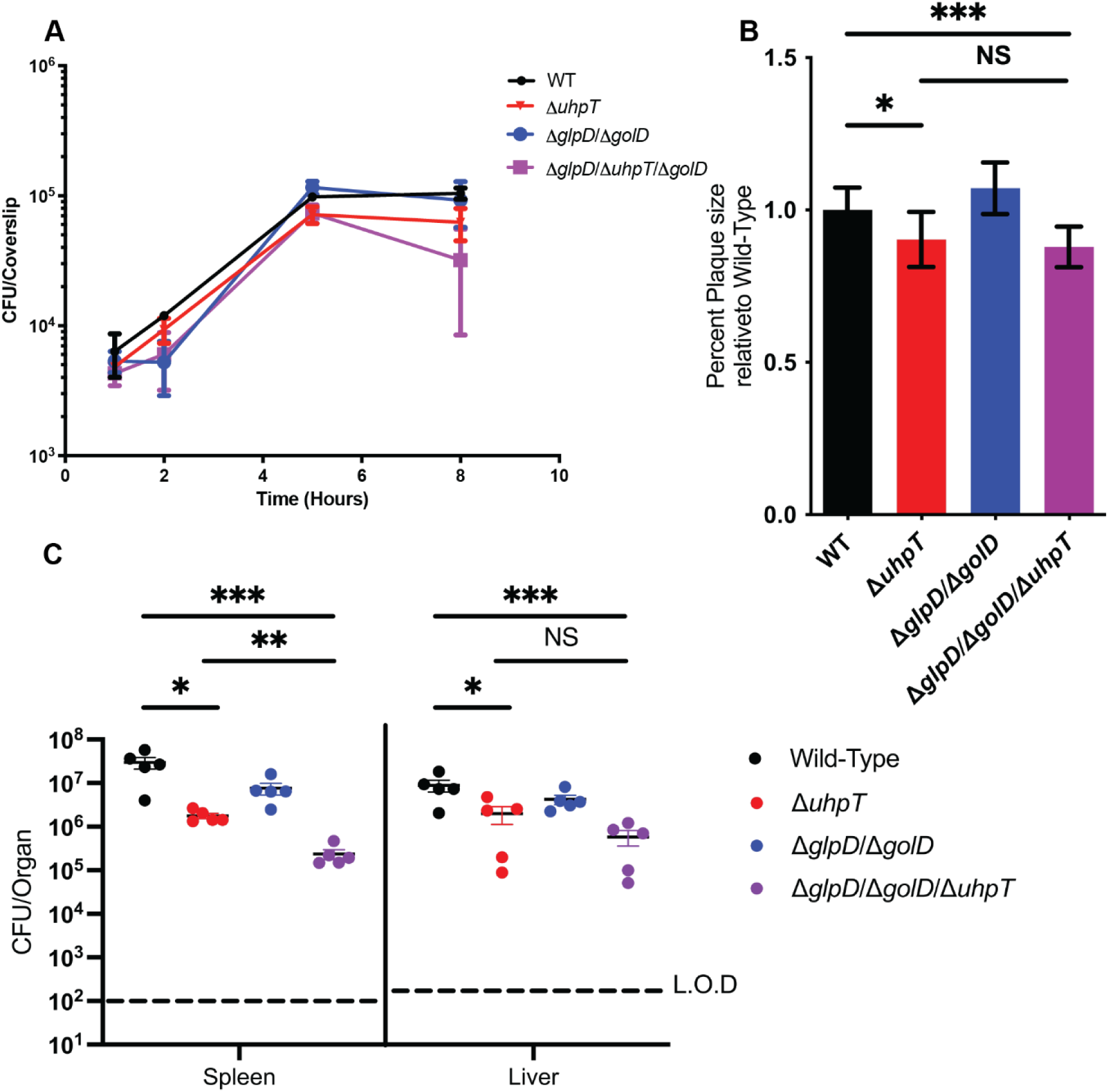
Mutants defective for metabolism of glycerol (Δ*glpD*/Δ*golD*) and/or hexose phosphates (Δ*uhpT*) are readily able to grow in the host cytosol and maintain virulence. (A) Intracellular growth of WT, Δ*glpD*/Δ*golD*, Δ*uhpT,* Δ*glpD*/Δ*golD*/Δ*uhpT* was determined in BMDMs following infection at an MOI of 0.2. Growth curves are representative of at least three independent experiments. Error bars represent the standard deviation of the means of technical triplicates within the representative experiment. (B) L2 fibroblasts were infected with indicated *L. monocytogenes* strains (MOI of 0.5) and were examined for plaque formation 4 days post infection. Data are normalized to WT plaque size and represent the standard deviation of the means from a single well’s plaques. (C) Bacterial burdens from the spleen and liver were enumerated at 48 hours post-intravenous infection with 1x10^5^ bacteria. Data are representative of results from two separate experiments. Horizontal dashed lines represent the limits of detection, and the bars associated with the individual strains represents the median and SEM of the group.

*In vivo*, *L. monocytogenes* not only replicates in the primary infected cell but must spread to neighboring cells using ActA mediated actin-based motility (22). This intracellular replication and spread is modeled *ex vivo* using a plaquing assay (58,59). Mutants with replication defects, impaired cytosolic survival, or defects in actin-based motility and secondary vacuole escape all produce small plaques as this assay measures both replication and cell-to-cell spread over multiple infectious cycles and days. Thus, plaque assays can further elucidate minor defects due to their more stringent conditions. To further test the hypothesis that glycerol and hexose phosphates contribute to cytosolic replication and cell-to-cell spread, we measured the plaque sizes of the metabolic mutants relative to WT *L. monocytogenes*. Hexose phosphate acquisition deficient *L. monocytogenes* (Δ*uhpT*) formed statistically significantly smaller plaques, albeit with a relatively modest defect (**Fig 2B**). In contrast, mutants defective for glycerol utilization (Δ*glpD*/Δ*golD*) generated plaques indistinguishable from WT. *L. monocytogenes*. Finally, a mutant defective for both glycerol and hexose phosphate utilization (Δ*glpD*/Δ*golD/*Δ*uhpT*) phenocopied a hexose phosphate mutant (Δ*uhpT*) alone suggesting no additive role for glycerol in the context of multicycle infection and cell to cell spread in fibroblasts (**Fig 2B**). This data indicates that while glycerol utilization is dispensable in single cycle infections in BMDMs and plaque multi-cycle plaque assays, hexose phosphates contribute to *L. monocytogenes* fitness in the context of multi-cycle infections and/or cell-to-cell spread, but not single cycle BMDM growth curves.

Given the apparent role for hexose phosphates in multicycle infections in the plaquing assay, we wanted to test the hypothesis that glycerol and hexose phosphates would be required for virulence *in vivo*. Mutants with replication and survival defects will sometimes show more robust virulence defects in murine models compared to *ex vivo* assays given the more restrictive physiology and multicycle infectious nature (60). To test this hypothesis, we utilized a murine disseminated listeriosis model and assessed virulence via bacterial burdens in the spleen and liver 48 hours after intravenous infection (**Fig 2C**). Consistent with results from the intracellular growth curves and the plaquing assay, glycerol mutants (Δ*glpD*/Δ*golD*) were not statistically significantly attenuated, although there was a trend towards lower bacterial burdens 48 hours post infection. Additionally, hexose phosphate mutants (Δ*uhpT*) showed mild but statistically significant attenuation in both organs with approximately 1.5-2 logs of virulence reduction.

Finally, failure to use glycerol in addition to hexose phosphates (Δ*glpD*/Δ*golD/*Δ*uhpT*) resulted in more significant attenuation than hexose phosphate mutants alone (Δ*uhpT*) in the spleen, while a similar trend was observed in the liver (**Fig 2C**). Taken together this data suggests that hexose phosphates, and not glycerol, are essential for multi-cycle replication *ex vivo* and full virulence during *in vivo* infection. Importantly, the relatively minor virulence defects for a mutant defective for utilization of both glycerol and hexose phosphates suggests that additional, yet to be defined carbon source(s) are the major contributors to replication and virulence of *L. monocytogenes in vivo*.

### BioLog phenotype microarray screening reveals WT and PrfA* *L. monocytogenes* equivalently respire on PTS mediated carbon sources

The lack of an intracellular growth defect of the Δ*glpD*/Δ*golD/*Δ*uhpT* mutant *ex vivo* and its relatively minor virulence defect *in vivo* led us to hypothesize that *L. monocytogenes* must use additional carbon sources during infection. To identify potential metabolites used by *L. monocytogenes* that could support cytosolic growth, we employed BioLog’s phenotypic carbon microarrays (PM1 and PM2A) to screen for differential carbon source respiration between WT and PrfA* *L. monocytogenes*. PrfA* mutants contain a G145S mutation in the virulence regulator PrfA that results in constitutive virulence gene expression and a physiologic state similar to that during infection, including upregulation of *uhpT* for the use of hexose phosphates (23,24). We hypothesized that PrfA*\* L. monocytogenes* may differentially use carbon sources relative to WT *L. monocytogenes*, similar to its differential use of hexose phosphates, which could reveal targets used to support cytosolic growth. Inoculation and setup of phenotypic microarray plates were performed as previously described, assays were performed in triplicate and plates were incubated at 37° stationary for 48 hours (61,62). At 48 hours post-inoculation, plates were assessed for change in tetrazolium dye color, indicating bacterial respiration on the carbon source. OD_490_ values were normalized to glucose, a carbon source known to be used by both strains, for each respective strain to account for potential global metabolic variance between strains. 190 total carbon sources were assessed for use by *L. monocytogenes* relative to glucose respiration (**Fig 3 and S4 Table**). WT *L. monocytogenes* was able to use 51 carbon sources at or above the level of glucose, of which, 35 are hypothesized to be available in the host cytosol according to the human metabolome database (63). PrfA* was able to consume all these same carbon sources and had significantly increased respiration on hexose phosphates as expected based on its known upregulation of *uhpT* (**Fig 3 and Table 1**). PrfA* also showed a statistically significant decreased respiration on a single PTS mediated carbon source, α-D-Lactose. Notably, PrfA* *L. monocytogenes* had overall similar use of most carbon sources, including PTS mediated carbon sources (**Fig 3 and S4 Table**). The glycerol and hexose phosphate (Δ*glpD*/Δ*golD*/Δ*uhpT*) mutant was similarly tested using PM1 and PM2A and phenocopied WT *L. monocytogenes* except for the loss of the ability to respire glycerol and α-Methyl-D-Glucoside (**S2 Fig and S1 Table 1**). Contrary to our hypothesis that PrfA* mutants would utilize metabolites differentially relative to WT *L. monocytogenes*, the only metabolites utilized more readily in PrfA* *L. monocytogenes* were hexose phosphates. Nevertheless, we were surprised to find that PrfA* *L. monocytogenes* readily utilized PTS-dependent carbon sources nearly identically to WT *L. monocytogenes.* This was striking given the established model in the field that PTS are not used during infection. However, PrfA* *L. monocytogenes* using PTS-mediated carbon sources is not mutually exclusive to the existing literature, in which, PTS transcripts are repressed during PrfA activation and use of PTS-mediated carbon sources’ reciprocally repress *prfA* expression (41,53).

**Figure 3.**
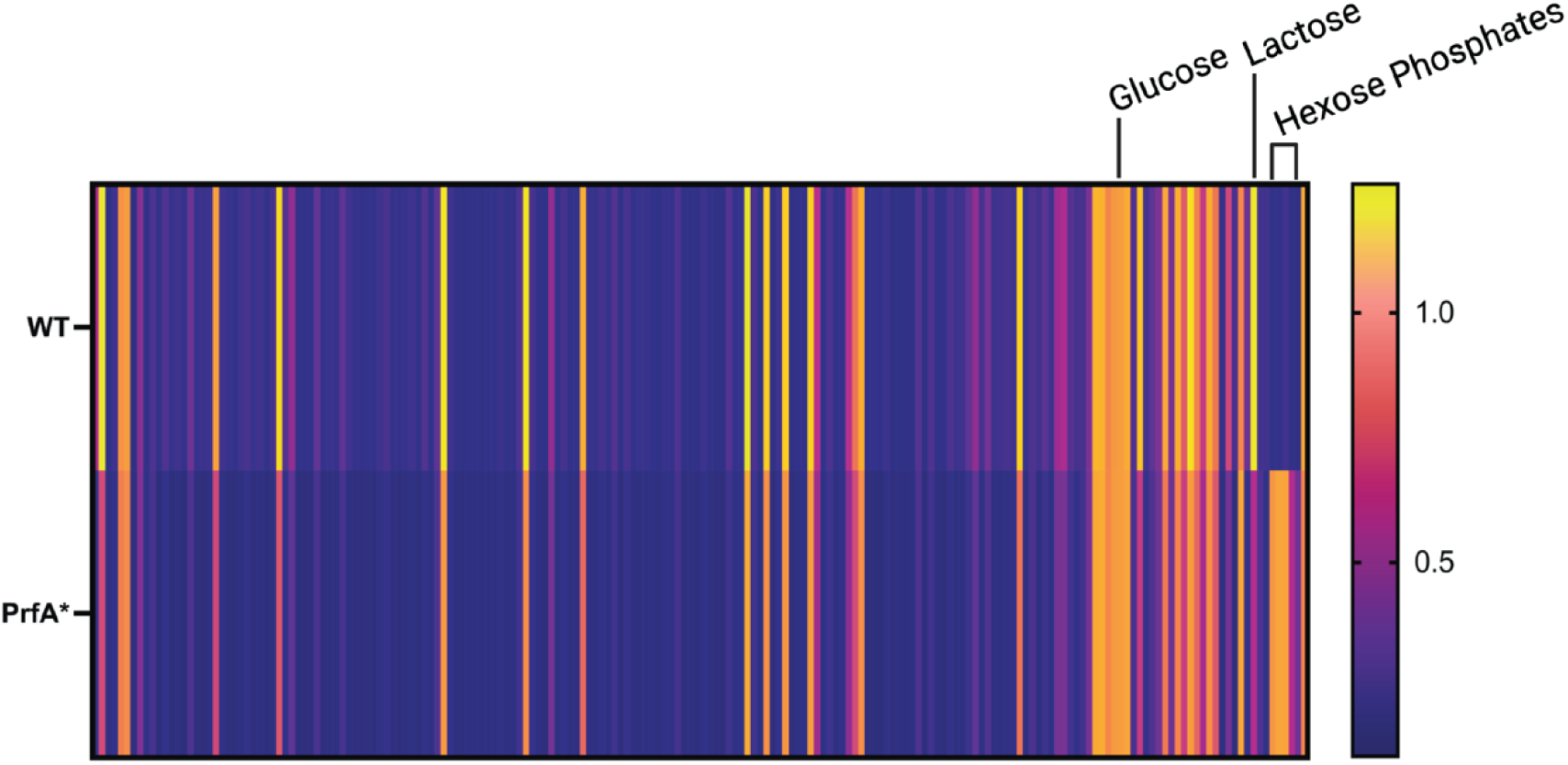
Carbon metabolite respiration of WT *and* PrfA* *L. monocytogenes* is globally similar, identified using Biolog’s Phenotypic Microarrays. Clustered heatmaps indicating level of tetrazolium dye color change as measured by OD_490_ at 48 hours in response to WT (Top) and PrfA* (Bottom) respiration of Carbon (PM1 & PM2A) at 37°C stationary. Each bar indicates the average of 3 biologic replicates. Samples were normalized to readings of a α-D-glucose control (∼1 on scale) and sorted based on cluster analysis.

**Table 1.** Statistically significant and greater than 2-fold differentially used metabolites between WT and PrfA* *L. monocytogenes*.

| Metabolite | Fold Change (WT/PrfA*) | P-Value |
| --- | --- | --- |
| D-Glucose-6-Phosphate | 0.16 | 0.000 |
| D-Glucose-1-Phosphate | 0.16 | 0.000 |
| D-Fructose-6-Phosphate | 0.21 | 0.000 |
| alpha-D-Lactose | 2.12 | 0.002 |

Purple highlights indicate metabolites used more in PrfA\* strains compared to wild-type and gold highlights are less used by PrfA\*.

### PTS are necessary for intracellular *L. monocytogenes* growth and virulence *in vivo*

Based on the similar utilization of PTS-mediated carbon sources between WT and PrfA* *L. monocytogenes* and the relatively minor virulence defects of the △*glpD*/△*golD*/△*uhpT* mutant, we hypothesized that PTS-dependent sugars could be an alternative carbon source during infection. The *L. monocytogenes* strain 10403s used in this study encodes 29 complete PTS systems encoded by 86 genes making it difficult to test individual PTS for importance during infection. Therefore, to test the hypothesis that PTS contribute to *L. monocytogenes* virulence we instead targeted the universally conserved phosphocarrier proteins essential for the function of all PTS, HPr (*ptsH*) and EI (*ptsI*) (**Fig 4A**). Δ*ptsI* mutants were constructed in WT and *△glpD/△golD/△uhpT L. monocytogenes* backgrounds to assess the relative contribution of hexose phosphates and glycerol vs PTS-dependent metabolites during cytosolic growth and virulence. To assess the role of PTS in cytosolic growth we infected primary murine BMDMs with WT, *△glpD/△golD/△uhpT,* Δ*ptsI*, *and △glpD/△golD/△uhpT/*Δ*ptsI L. monocytogenes*. While *△glpD/△golD/△uhpT* and WT *L. monocytogenes* were able to replicate in the macrophage cytosol as previously demonstrated (**Fig 2B and Fig 4B**), PTS deficient strains were completely unable to replicate, independent of the presence or absence of glycerol and hexose phosphate utilization pathways. Taken together these data demonstrate that PTS-dependent metabolites are both necessary and sufficient to support replication in the cytosol of BMDMs.

**Figure 4.**
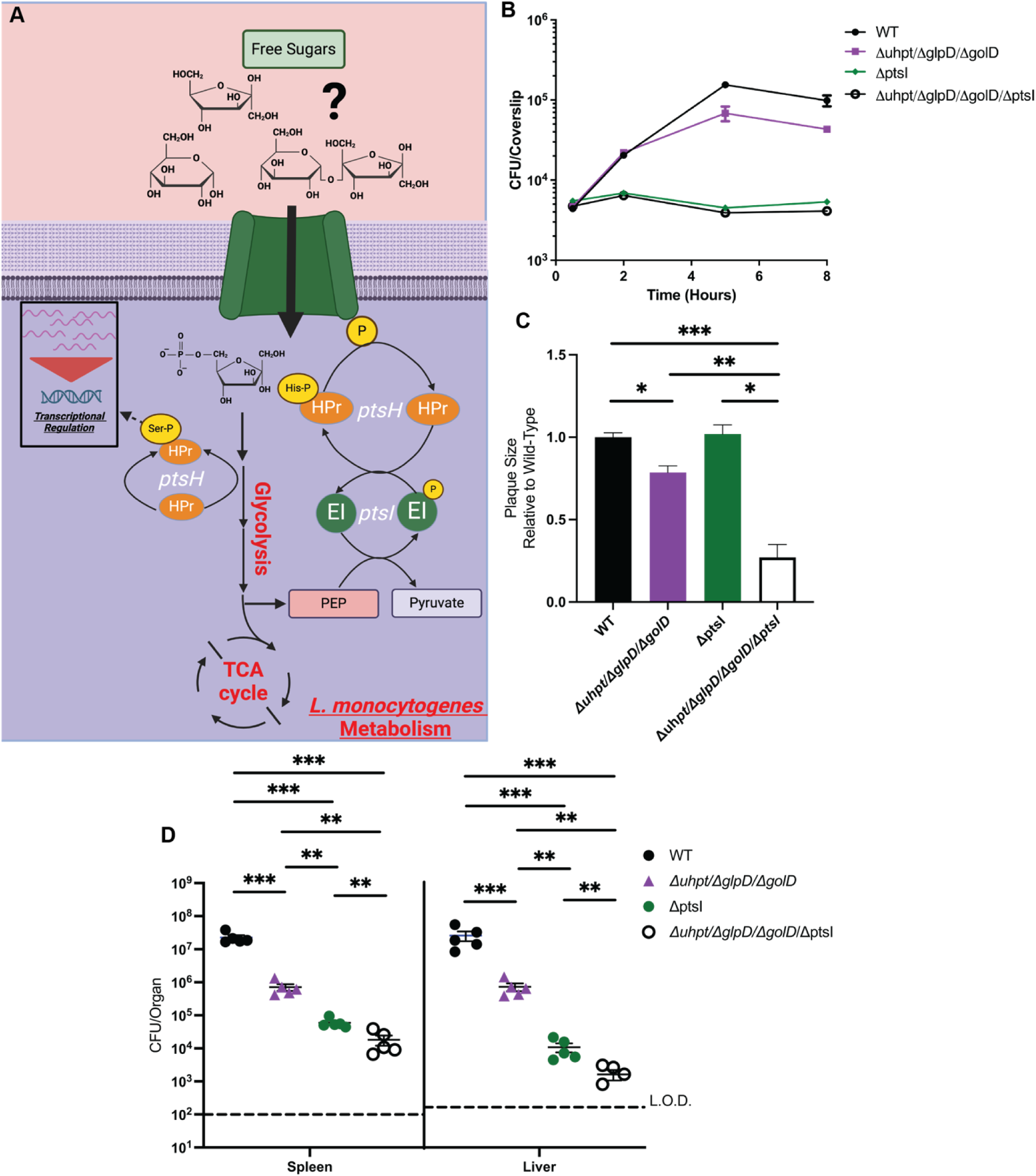
Δ*ptsI* mutants are impaired for intramacrophage growth and virulence, with more decreased virulence in a Δ*glpD*/Δ*golD*/Δ*uhpT* background. (A) PTS mediated free sugar import and phosphorylation by phosphocarrier protein phospho-cycling from the terminal conversion of phosphoenol-pyruvate (PEP) to pyruvate. (B) Intracellular growth of WT, Δ*glpD*/Δ*golD*/Δ*uhpT*, Δ*ptsI*, and Δ*glpD*/Δ*golD*/Δ*uhpT*/Δ*ptsI* was determined in BMDMs following infection at an MOI of 0.2. Growth curves are representative of at least three independent experiments. Error bars represent the standard deviation of the means of technical triplicates within the representative experiment. (C) L2 fibroblasts were infected with indicated *L. monocytogenes* strains (MOI of 0.5) and were examined for plaque formation 4 days post infection. Data are normalized to WT plaque size and represent the standard deviation of the means from a single well’s plaques. (D) Bacterial burdens from the spleen and liver were enumerated at 48 hours post-intravenous infection with 1x10^5^ bacteria. Data are representative of results from two separate experiments. Horizontal dashed lines represent the limits of detection, and the bars associated with the individual strains represents the median and SEM of the group.

The striking lack of cytosolic replication of PTS deficient *L. monocytogenes* in BMDMs, led us to ask whether PTS systems were similarly required for replication and cell-to-cell spread in more protracted multi-cycle infections by performing a plaquing assay in L2 fibroblasts. As previously demonstrated, *△glpD/△golD/△uhpT* mutants had a small plaquing defect relative to WT *L. monocytogenes*; however, in contrast to the BMDM growth phenotype of PTS deficient mutants, *L. monocytogenes* strains lacking only *ptsI* demonstrated no plaquing defects (**Fig 4C**).

Combining PTS deficiency with glycerol and hexose phosphate deficiency led to a significant decreased in plaque sizes (**Fig 4C**). Taken together these data suggest that while PTS are absolutely required for replication in the macrophage cytosol, PTS are only conditionally required in fibroblasts when glycerol and hexose phosphates are not available. These data suggest that there are differences in nutrient availability in different cell types, and further, that *L. monocytogenes* uses metabolic flexibility to grow in diverse cell types.

Given the differential requirements of PTS in macrophages versus fibroblasts *ex vivo*, we wanted to determine the role of PTS during *in vivo* infection in a murine disseminated listeriosis model. As we previously demonstrated, *△glpD/△golD/△uhpT* mutants had a minor, but statistically significant, virulence defect 48 hours post infection (**Fig 4D**). In contrast, Δ*ptsI* mutants were 500-fold and 5000-fold attenuated in the spleens and livers of infected mice respectively (**Fig 4D**). Furthermore, loss of PTS function in the absence of glycerol and hexose phosphate utilization led to an additive virulence defect whereby the *△glpD/△golD/△uhpT/*Δ*ptsI* mutants were more attenuated than the *△glpD/△golD/△uhpT* mutants or the Δ*ptsI* mutants alone (**Fig 4D**). Intra-macrophage growth, cell-to-cell spread, and virulence *in vivo* were proportionally rescued to genetic background levels with constitutive overexpression trans-complementation of *ptsI* (**S3 Fig)**. Together this data suggests that PTS mediated carbon acquisition is an important contributor to *in vivo* virulence. Furthermore, the additive attenuation of the *△glpD/△golD/△uhpT/*Δ*ptsI* mutant relative to *ΔptsI* mutant alone demonstrates that glycerol, hexose phosphates and PTS mediated carbon sources are all significant contributors to *L. monocytogenes* virulence *in vivo*.

To further confirm that the loss of virulence in the Δ*ptsI* mutant was due to loss of function of PTS and not another unknown function of EI, we created deletion mutants in the other conserved essential phosphocarrier protein HPr (*ptsH*). HPr is known to have a secondary role in genetic regulation via CcpA through phosphorylation by HPr kinase, therefore we hypothesized that Δ*ptsH* should, at minmum, phenocopy Δ*ptsI,* with the potential for further attenuation. Consistent with PTS being essential for growth in the macrophage cytosol, Δ*ptsH* mutants were unable to replicate in BMDM alone or when combined with hexose phosphate and glycerol (*△glpD/△golD/△uhpT*/Δ*ptsH*) mutants (**S4A Fig**). Similarly, Δ*ptsH* mutants were significantly attenuated for virulence *in vivo* and when combined with glycerol and hexose phosphate mutants, *△glpD/△golD/△uhpT*/Δ*ptsH* mutants were essentially avirulent *in vivo* (**S4B Fig)**. Taken together, these data suggest that PTS are major contributors to virulence of *L. monocytogenes in vivo* and that a loss of PTS combined with loss of glycerol and hexose phosphate utilization leads to almost complete attenuation of *L. monocytogenes in vivo*.

## DISCUSSION

Mechanisms of carbon acquisition, catabolism, and anabolism by cytosolic pathogens remain incompletely defined, but are vitally important virulence factors in driving pathogenesis. A mechanistic understanding of bacterial metabolism during infection can help identify novel anti-microbial targets and host targeted metabolic interventions. Despite *L. monocytogenes* being both an important pathogen and a model organism that has been studied for decades, we still have a relatively limited understanding of its metabolism during infection. In this work, we demonstrated that although hexose phosphate and glycerol do minimally contribute to *L. monocytogenes* infection *in vivo,* they are dispensable for cytosolic replication in macrophages in contrast to previous suggestions from isotopologue metabolomics. Further, combining an unbiased screen using BioLog’s Carbon Phenotypic Microarrays (PM1 and PM2A) with genetic deletions of conserved PTS phosphocarrier proteins, we demonstrate that PTS are essential for replication in the macrophage cytosol and are critical for virulence *in vivo*. Together our data suggest that *L. monocytogenes* is utilizing a previously underappreciated and more diverse metabolic strategy to replicate in the cytosolic environment and potentiate infection. These findings illuminate a generally overlooked contributor to virulence, PTS, and point to a system not previously identified as necessary for successful intracellular growth and virulence.

Despite PTS’ broad and well-defined role in carbon acquisition and previous demonstrated roles in the growth of *Shigella flexneri* and *Streptococcus pyogenes* during infection, they have largely been viewed as either minor contributors or in some cases active inhibitors of bacterial pathogenesis more broadly (39,41,43,51,53,64–66). In *L. monocytogenes,* work demonstrating that PTS activity represses *prfA* expression lead to a widespread and often repeated assumption that PTS are not used during infection, as virulence gene repression was considered detrimental to pathogenesis (38,43,44,55). Nevertheless, our findings demonstrate that a Δ*glpD*/Δ*golD/*Δ*uhpT* mutant *L. monocytogenes* with presumed obligate use of PTS, can still readily cause infection (31,53). However, it is possible that over reliance on PTS, and consequential PrfA regulon repression, resulted in the minor virulence defects of Δ*glpD*/Δ*golD/*Δ*uhpT L. monocytogenes* observed *in vivo*, while retaining intracellular growth. One remaining question is why *L. monocytogenes* would use this complex metabolic strategy utilizing such a wide array of carbon sources including those with potentially negative impacts on virulence gene expression? We hypothesize that *L. monocytogenes* may be using a balance of carbon sources to optimize virulence gene expression while maximizing carbon acquisition for metabolism and growth potential. As such, the phenotypes uncovered through these studies could represent both virulence gene dysregulation in addition to metabolic insufficiency due to failure of carbon source acquisition in the host

Bacterial PTS encode two phosphocarrier proteins that are necessary for the function of all PTS regardless of sugar: PtsI (EI) and PtsH (HPr) (42). Despite these two proteins sharing universal roles in PTS carbon acquisition, they have diverse functions, and per our results, phenotypes.

Δ*ptsI L. monocytogenes* are defective for only carbon acquisition through PTS as PtsI has no described role as a transcriptional regulator. In contrast, HPr (*ptsH*) has been shown previously to not only impact carbon acquisition through PTS but also to play a central role in transcriptional regulation of virulence through the phosphorylation of the HPr-Ser-46 residue (53,66). The HPr-Ser residue is phosphorylated by HPr Kinase (HprK) dependent on upper glycolytic flux and the abundance of fructose-1,6-bisphosphate and ATP. Importantly, these functions are not linked to lower glycolytic PEP to pyruvate conversion or PtsI (EI) function (65). The exact mechanism by which HPr-Ser-P enacts this repression is unknown (41,52).

Consistent with this, Δ*ptsH* (HPr) mutants in the WT and *△glpD/△golD/△uhpT L. monocytogenes* backgrounds show decreased virulence relative to those of Δ*ptsI* in an isogenic background suggesting that that the transcriptional dysregulation due to loss of *ptsH* is key to the additional virulence defects. Our favored hypothesis is that Δ*ptsH L. monocytogenes* is failing to modulate expression of genes necessary for alternate carbon source acquisition and/or deal with stresses unique to the host cytosol and these functions are retained in Δ*ptsI*. For example prior work has shown that HprK mediated phosphorylation of CcpA plays an important role in hierarchal carbon source utilization and stress responses in *L. monocytogenes* (67,68).

Because the exact transcriptional changes mediated by HPr-Ser-P are not well defined, it would be valuable to test how phospho-ablative and -mimetic HPr-Ser and HPr-His mutants behave differently through virulence assays and transcriptomic profiling. Overall, PtsI (EI) and PtsH (HPr) have separate but overlapping, functions that once characterized could unveil how and why bacteria connect the function of PTS to gene expression.

Previous isotopologue analysis by the Eisenreich and Goebel groups have shown that *L. monocytogenes* uses hexose phosphates and glycerol to support its cytosolic growth (28–32,69). These carbon substrates are logical carbon sources for an intracellular pathogen such as *L. monocytogenes* because not only does *L. monocytogenes* have dedicated transporters for these carbon sources but, equally importantly, they are available in the host cytosol (28,31). Nearly all sugar that is brought into eukaryotic cells is phosphorylated to prevent backward diffusion to the extracellular space. Once a sugar is phosphorylated it has two primary fates: 1. Funneling into glycolysis. 2. De-phosphorylation for use as a moiety/metabolite component in more complex forms, such as glycogen in the liver (70–72). Nevertheless, for PTS to be essential for *L. monocytogenes* growth in the cytosol, free sugars (monomers or polymers) must be present in abundance to support bacterial growth. Consistent with existing literature that PTS cannot facilitate transport of phosphorylated sugars, the *△glpD/△golD/△uhpT* mutant relying on PTS was unable to grow hexose phosphates *in vitro* but retained ability to grow in the host cytosol. Therefore, *L. monocytogenes* must have access to unphosphorylated sugar in the cytosol Whether these unphosphorylated, free sugars are liberated by the host or bacteria during the course of *L. monocytogenes* infection remains an unknown and critical component in understanding the use of PTS. One hypothesis is that *L. monocytogenes* is using a yet to be defined phosphatase to dephosphorylate and liberate free sugars. These putative phosphatases could represent high value targets for the development of antivirulence based antimicrobials (73) Alternatively, host cells could attempt to dephosphorylate sugars to allow sugar diffusion in an act of nutritional immunity, a process that *L. monocytogenes* might have evolved to take advantage of. Ultimately, understanding how *L. monocytogenes* accesses free sugars in the cytosol of an infected cell could inform whether other cytosolic pathogens could have access to and use similar carbon sources during infection.

Answering what specific PTS and corresponding carbon source might be used by *L. monocytogenes* is extremely challenging given the intertwined nature of the *L. monocytogenes*-macrophage metabolism and the redundancy of PTS systems in the *L. monocytogenes* genome (74) Importantly, while our work shows that the carbon acquisition function of PTS is an important contributor to *L. monocytogenes* cytosolic growth, we do not know specifically what PTS mediated sugars might be consumed. Because of the diverse arsenal of PTS encoded by *L. monocytogenes,* it is possible that there are multiple redundant carbon sources consumed by *L. monocytogenes* in the cytosol. One foothold to answer this question is the observation that *L. monocytogenes* PTS seem to have unique host cell and organ specific roles, indicating the presence of divergent metabolites within host cells that *L. monocytogenes* might be consuming. Identifying what specific carbon sources *L. monocytogenes* acquires from the host via PTS may unveil unique metabolic strategies to avoid host cell detection and support cytosolic bacterial physiology.

It is challenging to develop a complete picture of cytosolic pathogen metabolism given the ill-defined composition of the host cytosol, challenging technical methods, inherent metabolic redundancy, and diverse environments encountered in the host in different tissues. Our work demonstrates for the first time that PTS are essential for *L. monocytogenes* cytosolic growth and critical for virulence in contrast with the current dogma in the field (38,44,55). These highly conserved and redundant systems are understudied, and therefore their role in carbon acquisition by bacterial pathogens within the host cell is not well understood. Further work is needed to elucidate if PTS are critical for the cytosolic growth of other pathogens. Additionally, more work is needed to identify the preferred sugars for cytosolic pathogens and how these sugars are liberated. Finally, how differential carbon source availability across cell types, organ systems and even host species impacts bacterial pathogenesis remains to be elucidated.

## METHODS

### Ethics Statement

All animal-based experiments were performed using the protocol (M005916-R01-A01) approved by the Animal Use and Care Committee of the University of Wisconsin—Madison and consistent with the standards of the National Institutes of Health.

### Bacterial strains and culture

All *Listeria monocytogenes* strains used for experiments in this study were in a 10403s background. All *L. monocytogenes* strains were grown overnight in BHI and at 30°C stationary for all experiments, except as described. *Escherichia coli* strains were grown in Luria broth (LB) at 37°C shaking. Antibiotics used on *E. coli* were at a concentration of 100 µg/ml carbenicillin or 30 µg/ml kanamycin when appropriate. Antibiotics used on *L. monocytogenes* were at a concentration of 200 μg/mL streptomycin and/or 10 μg/mL chloramphenicol, when appropriate. Plasmids were transformed into chemically competent *E. coli* and further conjugated in *L. monocytogenes* using SM10 or S17 *E.coli*.

### Construction of strains

pLIM (from Arne Rietsche at Case Western) suicide plasmid was used for generation of in frame deletions (75). The pPL2 integrative vector pIMK2 was used for constitutive expression of *L. monocytogenes* genes (76). pLIM knockout constructs were cloned in XL1-Blue *E. coli* with 100 µg/ml carbenicillin (30μg/mL Kanamycin for pIMK2) and grown for plasmid harvest using Promega MIniPrep Kit. Harvested plasmid sequences were confirmed using was performed by Plasmidsaurus using Oxford Nanopore Technology with custom analysis and annotation.

Plasmid were then shuttled into *L. monocytogenes* through conjugation with SM10 (pLIM1) or S17 (pIMK2) *E. coli*. In-frame deletions of genes in *L. monocytogenes* were performed by allelic exchange using suicide plasmid pLIM as previously described with p-chlorophenylalanine as a counter selectable marker (77). Generated strains were frozen in 50:50 (glycerol:overnight culture) solution at -80°C.

### *In vitro* Growth Assays

Bacteria were grown overnight in BHI at 30°C stationary. Overnight cultures were used to generate inoculums with ∼3.7x10^7^ CFU/mL in PBS. 100 µLs per well of a flat bottom clear 96-well plate of *Listeria* synthetic media (LSM) with carbons source (supplied in 0.055 mM glucose carbon equivalents) was inoculated with 2 µL of inoculums. Plates were parafilmed on the edge to prevent evaporation and evaluated for OD_600_ in plate reader at 37°C shaking (250 r.p.m.) and reads every 15 minutes for times displayed.

### Intra-macrophage growth curves

Bone marrow-derived macrophages were isolated from CL57/BL6 mice and cultured as previously described (78). BMDMs were plated into 60 mm dishes contain 13 degassed coverslips. BMDMs cells were infected with *L. monocytogenes* strains at a multiplicity of infection [MOI] of 0.2. After 30 minutes BMDM media was exchanged for media containing 50 µg/ml Gentamycin. Coverslips were harvested, cells lysed in pure water, bacteria rescued isotonically, and plated to quantify CFU at displayed time points. All strains were assayed in biological triplicate and data displayed is one representative biologic replicate.

### Plaque Assay

Plaque assays were conducted using a L2 fibroblast cell line as previously described with minor modifications for visualization and quantification of plaques (58,59). L2 fibroblasts were seeded at 1.2 × 10^6^ per well of a 6-well plate, then infected at an MOI of 0.5 to obtain approximately 10-30 PFU per dish. At 4 days postinfection, cells were stained with 0.3% crystal violet for 10 min and washed twice with deionized water. Stained wells were scanned, uploaded, and areas of plaque formation were measured on ImageJ analysis software. All strains were assayed in biological triplicate and the plaque areas of each strain were normalized to wild-type plaque size within each replicate.

### Murine Infection and Organ Burdens

Infections were performed as previously described (79). Briefly, 6 to 8-week-old female and male C57BL/6 mice were infected IV with 1×10^5^ CFU logarithmically growing *L. monocytogenes* (optical density at 600 nm [OD600] = 0.5) in 200 µL of PBS. 48 hours post-infection, mice were euthanized, and livers and spleens were harvested, homogenized in water with 0.1% NP-40, and plated for CFU. Two independent replicates of each experiment with 5 mice per group were performed.

### Cell Culture

L2 cells were all kind gifts from Daniel Portnoy (UC Berkeley) (59). Bone marrow-derived macrophages (BMDM) were prepared from 6-to-8-week-old mice as previously described (78).

### Phenotypic Microarrays

Phenotype Microarrays 1 (Cat. #12111) and 2A (Cat. #12112) were obtained from BioLog (BioLog). Plates were prepared and inoculated as previously described (62,80). OmniLog incubation was substituted with incubation at 37°C stationary. OD490 was collected for each plate at 24 and 48 hours. Data was then normalized to consumption of α-D-glucose for each strain and replicate, and averaged across triplicate. Value were clustered based on similarity using clustergrammer and plotted as a heat-map in Prism 6 (81). Data normalized to glucose was used for statistical analysis and displayed in tables. Statistics are representative of a student’s T-test between two strains.

### Statistical Analysis

Prism 6 (GraphPad Software) was used for statistical analysis of data. Means from two groups of BioLog plates were compared with unpaired two-tailed Student’s T-test. Means from more than two groups for all other assays were analyzed by Kruskal-Wallis test. Independently, Mann-Whitney Test was used to analyze two group comparison of non-normal data from animal experiments. * *p* < 0.05, ** *p* < 0.01, *** *p* < 0.001 for all statistical tests displayed.

---

**Supplemental Figure 1.**
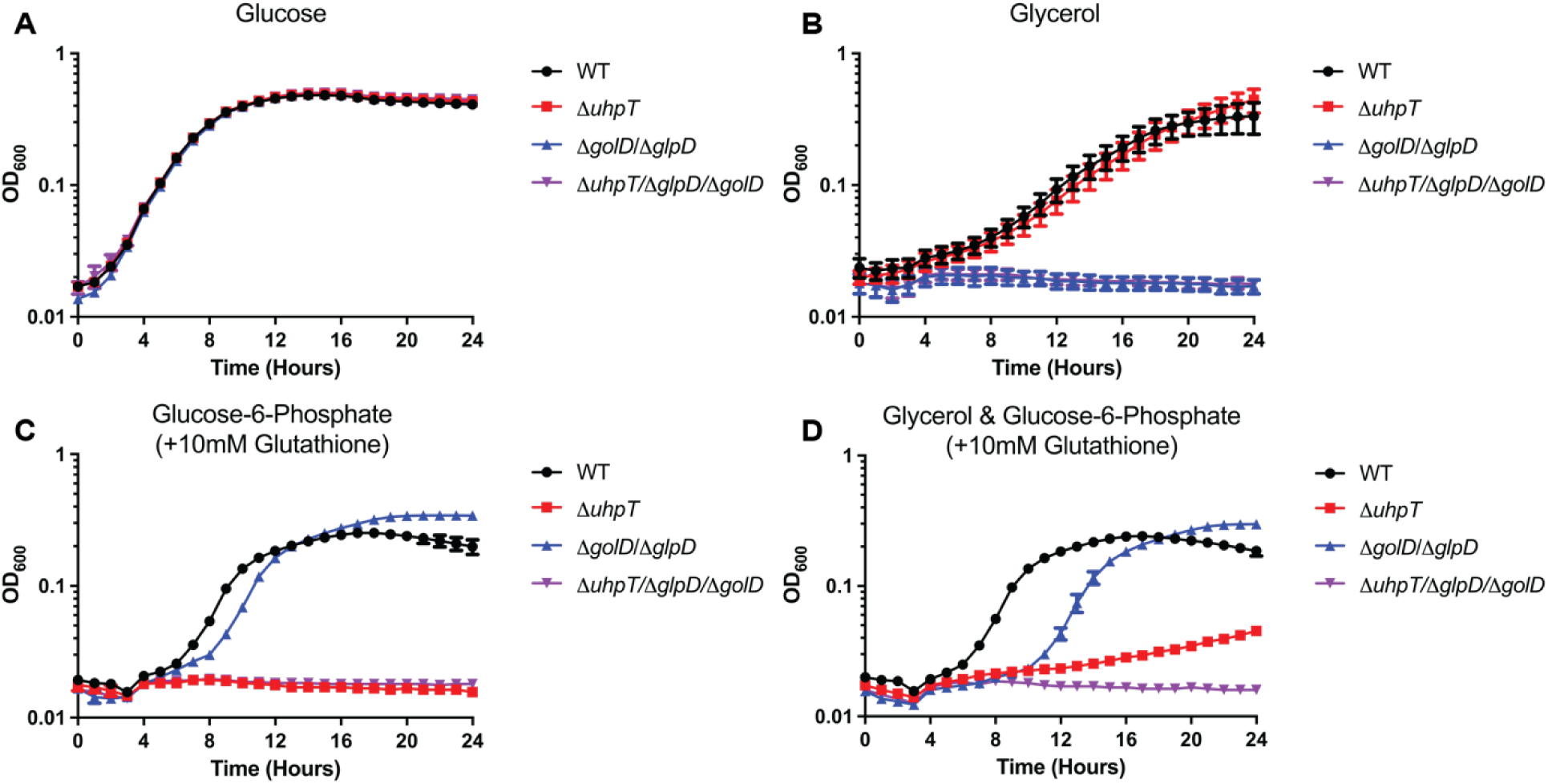
Mutants defective for consumption of glycerol (Δ*glpD*/Δ*golD)* and/or hexose phosphates (Δ*uhpT*) require alternative carbon sources to support growth in a defined medium. Indicated strains were grown in LSM at 37°C, shaking at 250 r.p.m. with the addition of 55mM glucose (A) or carbon equivalent amounts of hexose phosphates (10mM glutathione) and glycerol (B). OD_600_ was monitored every 15 minutes for 24 hours. Data represents average of three technical replicates from one representative of three biological replicates.

**Supplemental Figure 2.**
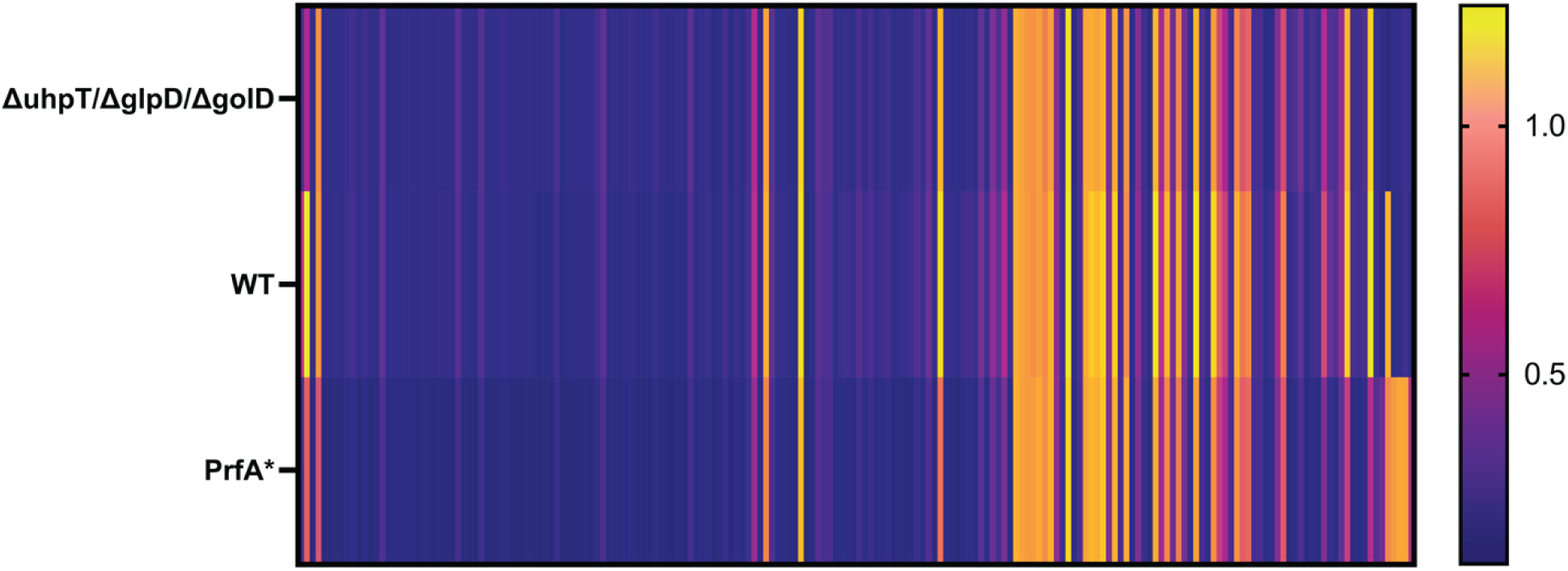
A △*glpD*/△*golD*/△*uhpT mutant* shows near identical respiration of carbon metabolites to that of WT *L. monocytogenes.* Clustered heatmaps indicating level of tetrazolium dye color change as measured by OD_490_ at 48 hours in response to △*glpD*/△*golD*/△*uhpT* (Top), WT (Middle), and PrfA* (Bottom) respiration of Carbon (PM1 & PM2) at 37°C stationary. Each bar indicates the average of 3 biologic replicates. Samples were normalized to readings of a α-D-glucose control (∼1 on scale) and sorted based on cluster analysis.

**Supplemental Figure 3.**
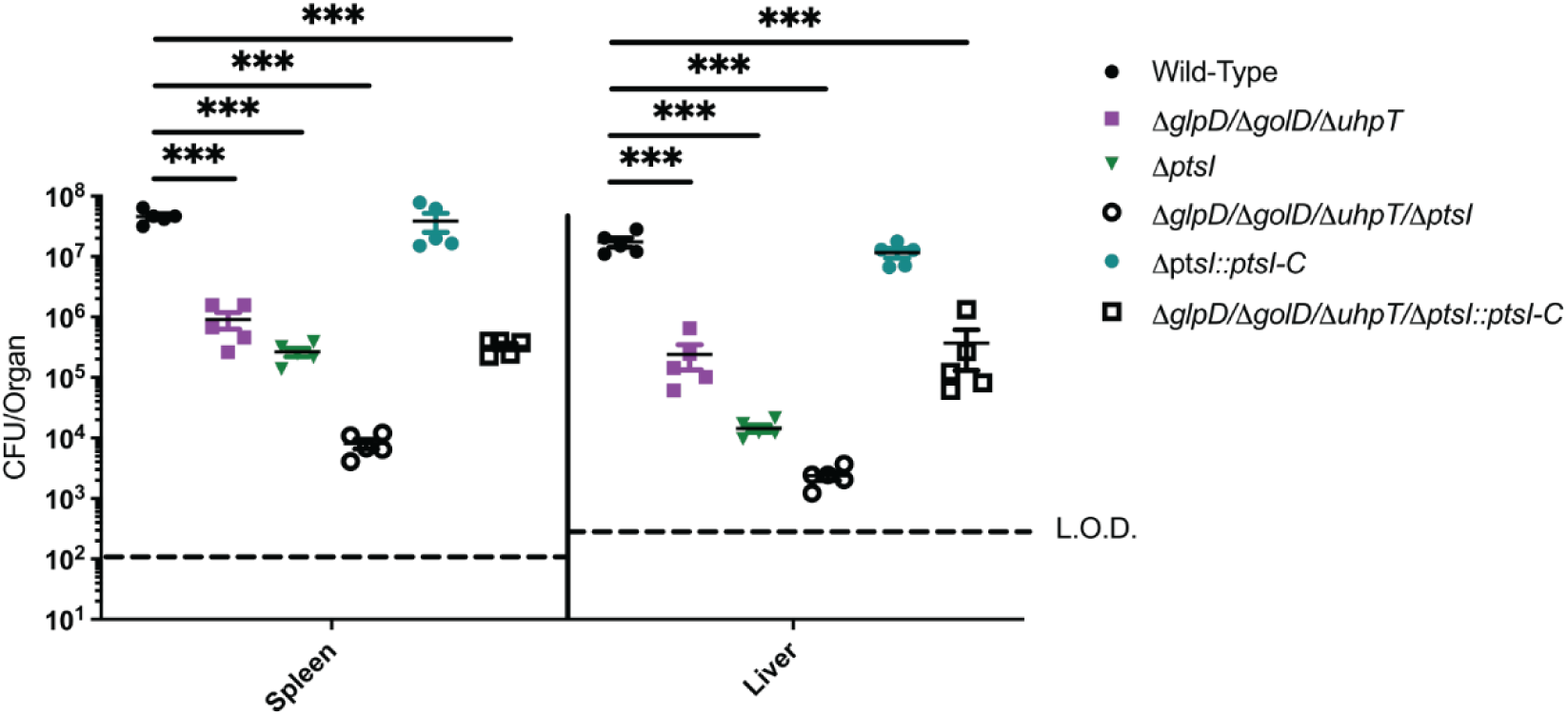
Δ*p*tsI mutants can be trans-complemented with *ptsI* over expression for intracellular growth and virulence. Bacterial burdens from the spleen and liver were enumerated at 48 hours post-intravenous infection with 1x10^5^ bacteria. Data are representative of results from two separate experiments. Horizontal dashed lines represent the limits of detection, and the bars associated with the individual strains represents the median and SEM of the group.

**Supplemental Figure 4.**
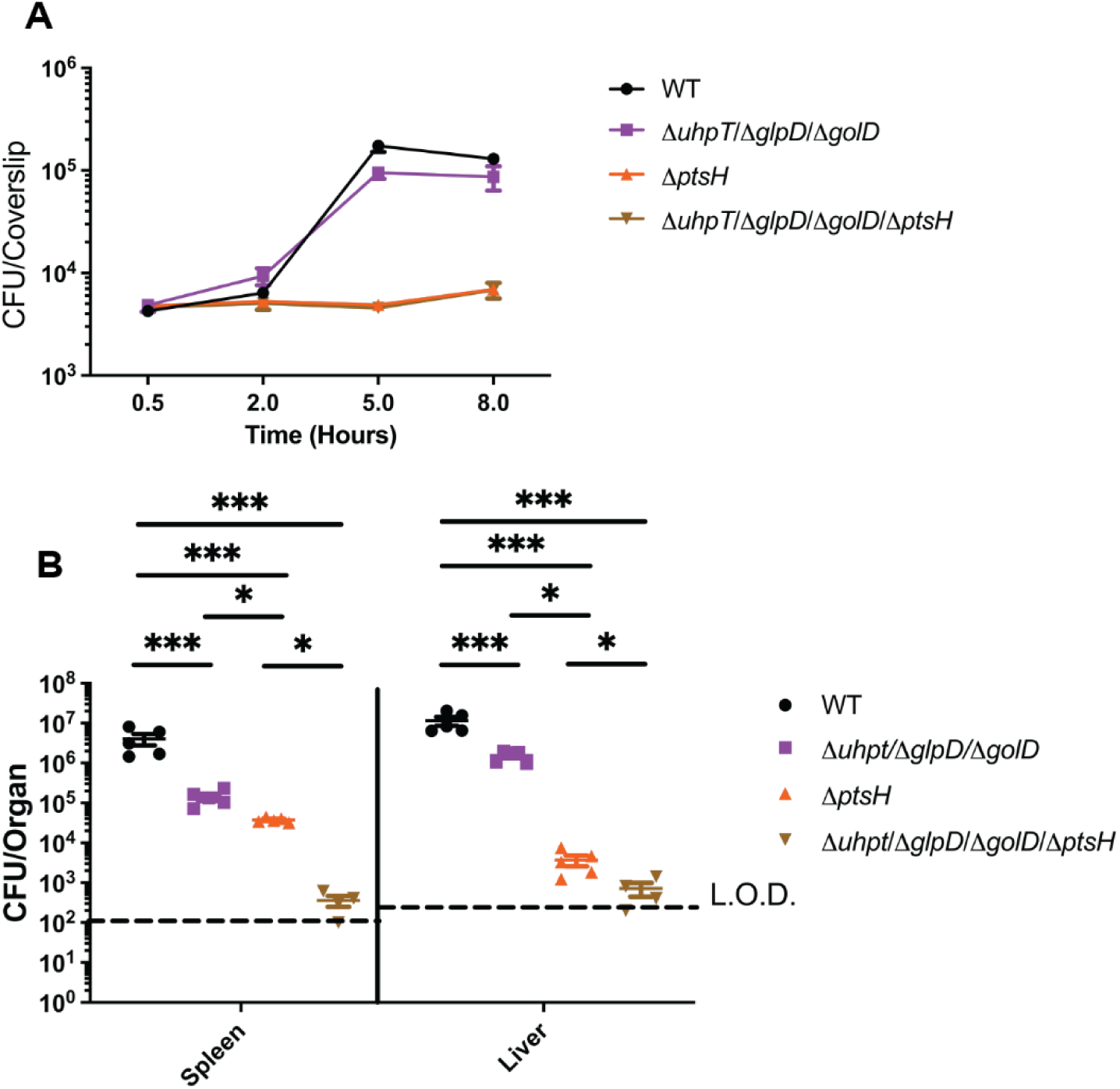
Δ*p*tsH mutants are more attenuated for intracellular growth and virulence than in Δ*ptsI* in all backgrounds. (A) Intracellular growth of wild-type, Δ*glpD*/Δ*golD*/Δ*uhpT*, Δ*ptsH*, and Δ*glpD*/Δ*golD*/Δ*uhpT*/Δ*ptsH* was determined in BMDMs following infection at an MOI of 0.2. Growth curves are representative of at least three independent experiments. Error bars represent the standard deviation of the means of technical triplicates within the representative experiment. (B) Bacterial burdens from the spleen and liver were enumerated at 48 hours post-intravenous infection with 1x10^5^ bacteria. Data are representative of results from two separate experiments. Horizontal dashed lines represent the limits of detection, and the bars associated with the individual strains represents the median and SEM of the group.

**Supplemental Table 1.** Statistically significant and greater than 2-fold differentially used metabolites between WT and △*glpD*/△*golD*/△*uhpT L. monocytogenes*.

| Metabolite | Fold Change<br>(WT/$\Delta glpD/\Delta golD/\Delta uhpT$) | P-Value |
| --- | --- | --- |
| Glycerol | 7.97 | 0.001 |
| alpha-Methyl-D-Glucoside | 2.25 | 0.005 |

Purple highlights indicate metabolites used more in WT strains compared to $\Delta glpD/\Delta golD/\Delta uhpT$ *L. monocytogenes*.

**Supplemental Table 2.** Bacterial Strains Used.

| Strain | Description | Reference |
| --- | --- | --- |
| XL1-Blue | competent *E. coli* strain | (60) |
| SM10 | *E. coli* strain for conjugations into *L. monocytogenes*; Km<sup>R</sup> | (60) |
| S17 | *E. coli* strain for conjugations into *L. monocytogenes*; Sp<sup>R</sup> | (76) |
| 10403S [JDS1] | Background *L. monocytogenes* 10403s strain | (82) |
| MJF35 | $\Delta uhpT$ | This Work & (28) |
| MJF112 | $\Delta glpD/\Delta golD$ | This Work |
| MJF121 | $\Delta glpD/\Delta golD/\Delta uhpT$ | This Work |
| MJF279 | $\Delta ptsI$ | This Work |
| MJF281 | $\Delta glpD/\Delta golD/\Delta uhpT/\Delta ptsI$ | This Work |
| MJF286 | $\Delta ptsI::ptsI-C$ (pIMK2) | This Work |
| MJF288 | $\Delta glpD/\Delta golD/\Delta uhpT/\Delta ptsI::ptsI-C$ (pIMK2) | This Work |
| MJF223 | $\Delta ptsH$ | This Work & (53) |
| MJF225 | $\Delta glpD/\Delta golD/\Delta uhpT/\Delta ptsH$ | This Work |

**Supplemental Table 3.**
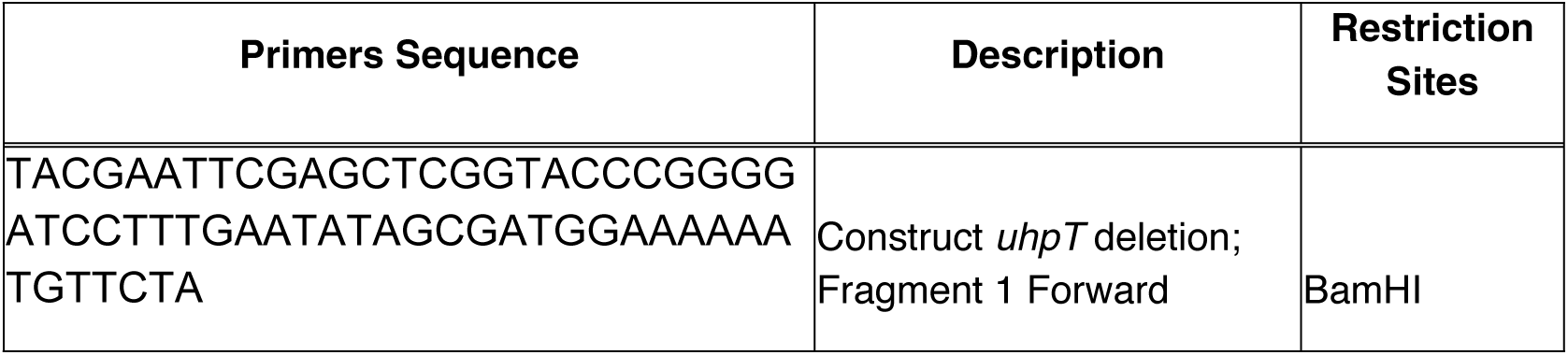
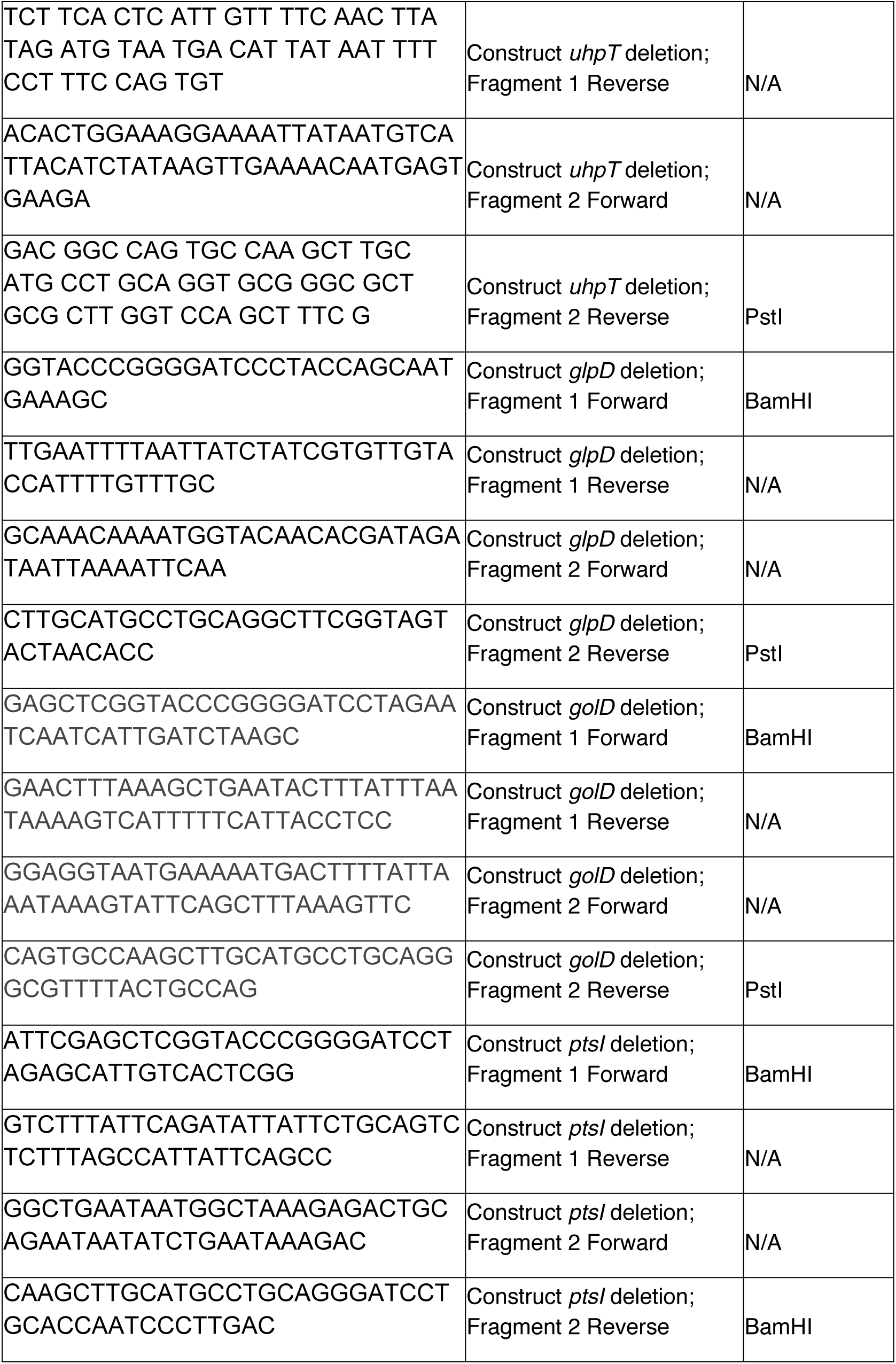
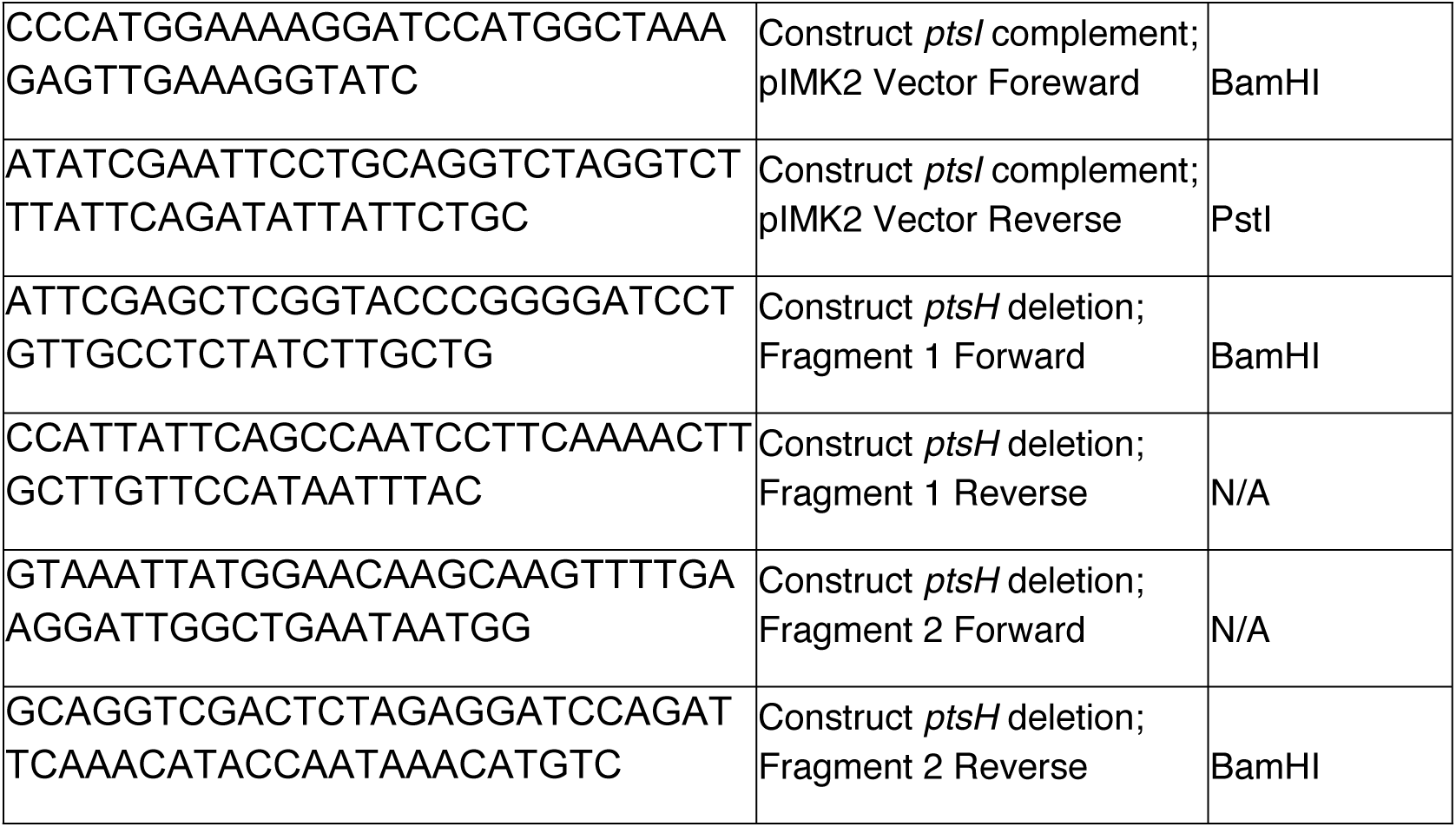
Primers Used.

**Supplemental Table 4.** Results of BioLog PM1 and PM2A at 48 hours for WT, Δ*glpD*/Δ*golD*/Δ*uhpT*, and PrfA*.

| <u>Metabolite</u> | <u>Fold Usage Normalized to α-D-Glucose</u> |  |  | <u>P-value (Student's T-Test)</u> |  |
| --- | --- | --- | --- | --- | --- |
|  | <u>WT</u> | <u>*ΔglpD/ΔgldI/ΔuhpT*</u> | <u>PrfA*</u> | <u>WT v.s. *ΔglpD/ΔgldI/ΔuhpT*</u> | <u>WT v.s. PrfA*</u> |
| Amygdalin | 1.24 | 1.18 | 1.15 | 0.25 | 0.10 |
| N-Acetyl-D-Glucosamine | 1.06 | 0.99 | 1.05 | 0.30 | 0.76 |
| Maltotriose | 1.08 | 1.01 | 1.04 | 0.22 | 0.18 |
| D-Trehalose | 0.93 | 0.96 | 1.03 | 0.39 | 0.01 |
| D-Glucose-6-Phosphate | 0.17 | 0.17 | 1.02 | 0.97 | 0.00 |
| D-Fructose-6-Phosphate | 0.21 | 0.19 | 1.01 | 0.68 | 0.00 |
| Beta-Methyl-D-Glucoside | 1.03 | 1.02 | 1.01 | 0.78 | 0.72 |
| alpha-Cyclodextrin | 1.14 | 1.05 | 1.11 | 0.41 | 0.71 |
| alpha-D-Glucose | 1.00 | 1.00 | 1.00 | N/A | N/A |
| D-Mannose | 1.18 | 1.06 | 0.99 | 0.14 | 0.01 |
| D-Cellobiose | 0.98 | 0.96 | 0.99 | 0.70 | 0.83 |
| Arbutin | 1.17 | 1.15 | 1.08 | 0.53 | 0.06 |
| D-Glucose-1-Phosphate | 0.16 | 0.17 | 0.98 | 0.58 | 0.00 |
| Beta-Cyclodextrin | 1.12 | 1.04 | 1.07 | 0.24 | 0.30 |
| D-Fructose | 1.06 | 0.91 | 0.96 | 0.25 | 0.38 |
| Beta-D-Allose | 1.12 | 1.04 | 1.04 | 0.25 | 0.16 |
| Xylitol | 1.18 | 1.06 | 1.03 | 0.30 | 0.24 |
| Sailcin | 1.03 | 1.03 | 1.05 | 0.98 | 0.63 |
| Gamma-Cyclodextrin | 0.98 | 0.99 | 1.02 | 0.88 | 0.55 |
| N-Acetyl-Beta-D-Mannosamine | 0.92 | 0.88 | 0.93 | 0.23 | 0.82 |
| Glycerol | 1.05 | 0.13 | 0.92 | 0.00 | 0.31 |
| L-Rhamnose | 1.07 | 0.94 | 0.92 | 0.35 | 0.32 |
| Gentiobiose | 0.94 | 0.96 | 0.99 | 0.55 | 0.05 |
| D-Psicose | 1.05 | 0.92 | 0.89 | 0.31 | 0.20 |
| Dihydroxy Acetone | 1.15 | 0.99 | 0.94 | 0.24 | 0.14 |
| D-Glucosamine | 1.04 | 1.00 | 0.91 | 0.71 | 0.12 |
| Uridine | 0.93 | 0.80 | 0.81 | 0.45 | 0.48 |
| Alpha-Methyl-D-Mannoside | 1.17 | 1.08 | 0.86 | 0.36 | 0.12 |
| L-Lyxose | 0.89 | 0.70 | 0.75 | 0.21 | 0.36 |
| 2-Deoxy-D-Ribose | 0.86 | 0.74 | 0.77 | 0.34 | 0.44 |
| alpha-Methyl-D-Glucoside | 1.23 | 0.55 | 0.79 | 0.00 | 0.02 |
| D-Arabitol | 0.99 | 0.95 | 0.74 | 0.77 | 0.01 |
| D-Ribose | 0.78 | 0.68 | 0.65 | 0.41 | 0.27 |
| Maltose | 1.12 | 1.07 | 0.64 | 0.25 | 0.00 |
| alpha-D-Lactose | 1.24 | 1.14 | 0.59 | 0.30 | 0.00 |
| D-Malic Acid | 0.14 | 0.16 | 0.58 | 0.09 | 0.39 |
| D-Xylose | 0.65 | 0.54 | 0.54 | 0.47 | 0.49 |
| Acetic Acid | 0.59 | 0.57 | 0.51 | 0.69 | 0.35 |
| 5-Keto-D-Gluconic Acid | 0.60 | 0.53 | 0.52 | 0.27 | 0.20 |
| Capric Acid | 0.46 | 0.43 | 0.46 | 0.85 | 0.98 |
| Inosine | 0.57 | 0.48 | 0.40 | 0.51 | 0.23 |
| N-Acetyl-D-Glucosaminitol | 0.53 | 0.43 | 0.43 | 0.27 | 0.25 |
| Acetoacetic Acid | 0.41 | 0.37 | 0.38 | 0.73 | 0.80 |
| Sorbic Acid | 0.48 | 0.44 | 0.42 | 0.39 | 0.23 |
| Dextrin | 0.70 | 0.57 | 0.42 | 0.33 | 0.07 |
| L-Arabinose | 0.42 | 0.37 | 0.36 | 0.51 | 0.47 |
| m-Tartaric Acid | 0.14 | 0.16 | 0.35 | 0.08 | 0.42 |
| Adenosine | 0.40 | 0.35 | 0.34 | 0.57 | 0.48 |
| D-Arabinose | 0.48 | 0.40 | 0.34 | 0.21 | 0.04 |
| alpha-keto-Butyric Acid | 0.48 | 0.48 | 0.30 | 1.00 | 0.00 |
| Palatinose | 0.38 | 0.33 | 0.28 | 0.25 | 0.02 |
| Mucic Acid | 0.31 | 0.39 | 0.25 | 0.50 | 0.59 |
| D-Fucose | 0.28 | 0.30 | 0.27 | 0.69 | 0.78 |
| Beta-Methyl-D-Galactoside | 0.30 | 0.28 | 0.26 | 0.81 | 0.54 |
| Putrescine | 0.20 | 0.18 | 0.27 | 0.71 | 0.28 |
| D-Tagatose | 0.36 | 0.32 | 0.26 | 0.59 | 0.18 |
| Thymidine | 0.24 | 0.27 | 0.23 | 0.50 | 0.73 |
| 3-Methyl Glucose | 0.32 | 0.31 | 0.25 | 0.94 | 0.50 |
| Adonitol | 0.25 | 0.23 | 0.22 | 0.70 | 0.66 |
| Oxalic Acid | 0.29 | 0.31 | 0.24 | 0.79 | 0.22 |
| Glyoxylic Acid | 0.24 | 0.26 | 0.22 | 0.88 | 0.65 |
| Sedoheptulosan | 0.25 | 0.24 | 0.24 | 0.88 | 0.84 |
| Glucuronamide | 0.27 | 0.24 | 0.21 | 0.15 | 0.03 |
| D,L-Octopamine | 0.22 | 0.20 | 0.23 | 0.65 | 0.90 |
| 2-Deoxy-Adenosine | 0.26 | 0.29 | 0.20 | 0.56 | 0.29 |
| Pyruvic Acid | 0.26 | 0.27 | 0.20 | 0.69 | 0.01 |
| Oxalomalic Acid | 0.27 | 0.24 | 0.22 | 0.59 | 0.31 |
| 2,3-Butanedione | 0.25 | 0.22 | 0.22 | 0.38 | 0.37 |
| Mannan | 0.24 | 0.21 | 0.22 | 0.57 | 0.67 |
| L-Histidine | 0.23 | 0.21 | 0.21 | 0.57 | 0.41 |
| Turanose | 0.31 | 0.25 | 0.21 | 0.30 | 0.09 |
| Pectin | 0.29 | 0.29 | 0.21 | 0.86 | 0.06 |
| Sebacic Acid | 0.60 | 0.22 | 0.20 | 0.21 | 0.19 |
| L-Sorbose | 0.23 | 0.22 | 0.20 | 0.88 | 0.56 |
| alpha-Keto-Valeric Acid | 0.24 | 0.21 | 0.19 | 0.60 | 0.35 |
| Sec-Butylamine | 0.22 | 0.22 | 0.19 | 0.94 | 0.37 |
| Beta-Methyl-D-Xyloside | 0.21 | 0.24 | 0.19 | 0.54 | 0.66 |
| L-Fucose | 0.22 | 0.21 | 0.17 | 0.80 | 0.27 |
| N-Acetyl-Neuaminic Acid | 0.21 | 0.20 | 0.19 | 0.48 | 0.31 |
| Methyl Pyruvate | 0.20 | 0.21 | 0.17 | 0.46 | 0.02 |
| Propionic Acid | 0.20 | 0.24 | 0.17 | 0.32 | 0.20 |
| 3-0-Beta-D-Galactopyranosyl-D-Arabinose | 0.26 | 0.24 | 0.18 | 0.67 | 0.07 |
| L-Alanyl-Glycine | 0.25 | 0.26 | 0.16 | 0.34 | 0.00 |
| L-Galactonic Acide-gamma-lactone | 0.19 | 0.19 | 0.16 | 0.80 | 0.23 |
| Inulin | 0.34 | 0.27 | 0.18 | 0.46 | 0.11 |
| L-Pyrogutamic Acid | 0.21 | 0.19 | 0.17 | 0.49 | 0.30 |
| D-Melezitose | 0.23 | 0.21 | 0.18 | 0.67 | 0.31 |
| L-Lactic Acid | 0.26 | 0.25 | 0.16 | 0.83 | 0.03 |
| L-Alanine | 0.19 | 0.21 | 0.16 | 0.59 | 0.26 |
| Tween 40 | 0.18 | 0.19 | 0.15 | 0.51 | 0.14 |
| D-Sorbitol | 0.21 | 0.18 | 0.15 | 0.08 | 0.02 |
| D-Galactose | 0.20 | 0.17 | 0.15 | 0.57 | 0.29 |
| D,L-alpha-glycerol-phosphate | 0.17 | 0.17 | 0.15 | 0.85 | 0.38 |
| D-Galactonic Acid-gamma-Lactone | 0.16 | 0.16 | 0.15 | 0.55 | 0.57 |
| alpha-Methyl-D-Galactoside | 0.18 | 0.18 | 0.15 | 0.99 | 0.49 |
| D-Glucuronic Acid | 0.19 | 0.19 | 0.15 | 0.89 | 0.20 |
| L-Alaninamide | 0.20 | 0.19 | 0.16 | 0.94 | 0.23 |
| Butyric Acid | 0.20 | 0.26 | 0.16 | 0.47 | 0.19 |
| 2-Aminoethanol | 0.17 | 0.18 | 0.14 | 0.44 | 0.14 |
| L-Methionine | 0.20 | 0.16 | 0.16 | 0.46 | 0.55 |
| alpha-Hydroxy Butyric Acid | 0.21 | 0.21 | 0.14 | 0.96 | 0.13 |
| D-Galacturonic Acid | 0.18 | 0.18 | 0.14 | 0.93 | 0.04 |
| L-Arabitol | 0.19 | 0.16 | 0.16 | 0.34 | 0.26 |
| Maltitol | 0.22 | 0.18 | 0.16 | 0.47 | 0.29 |
| D-Mannitol | 0.17 | 0.17 | 0.14 | 0.83 | 0.20 |
| Tween 80 | 0.17 | 0.17 | 0.14 | 1.00 | 0.09 |
| m-Hydroxy Phenyl Acetic Acid | 0.17 | 0.18 | 0.14 | 0.57 | 0.06 |
| Glycogen | 0.21 | 0.19 | 0.15 | 0.49 | 0.09 |
| Dulcitol | 0.17 | 0.16 | 0.14 | 0.69 | 0.19 |
| Glycyl-L-Proline | 0.22 | 0.19 | 0.14 | 0.17 | 0.02 |
| Beta-Methyl-D-Glucuronic Acid | 0.20 | 0.18 | 0.15 | 0.64 | 0.24 |
| Lactitol | 0.20 | 0.19 | 0.15 | 0.81 | 0.26 |
| Delta-Amino Valeric Acid | 0.16 | 0.15 | 0.15 | 0.70 | 0.75 |
| L-Glucose | 0.20 | 0.18 | 0.15 | 0.58 | 0.23 |
| L-Leucine | 0.16 | 0.15 | 0.15 | 0.55 | 0.62 |
| Glycine | 0.18 | 0.17 | 0.15 | 0.68 | 0.27 |
| Caproic Acid | 0.18 | 0.17 | 0.15 | 0.58 | 0.20 |
| L-Proline | 0.18 | 0.18 | 0.14 | 0.94 | 0.07 |
| Succinic Acid | 0.17 | 0.18 | 0.14 | 0.72 | 0.06 |
| D-Gluconic Acid | 0.17 | 0.18 | 0.14 | 0.61 | 0.12 |
| Negative Control | 0.18 | 0.17 | 0.14 | 0.84 | 0.03 |
| Negative Control | 0.18 | 0.16 | 0.15 | 0.49 | 0.43 |
| 3-Hydroxy-2-Butanone | 0.21 | 0.17 | 0.15 | 0.41 | 0.27 |
| Bromo Succinic Acid | 0.15 | 0.18 | 0.13 | 0.35 | 0.21 |
| D-Alanine | 0.16 | 0.17 | 0.13 | 0.55 | 0.21 |
| 2-Hydroxy Benzoic Acid | 0.16 | 0.17 | 0.15 | 0.62 | 0.39 |
| L-Aspartic Acid | 0.17 | 0.16 | 0.13 | 0.50 | 0.03 |
| Apha-Hydroxy Glutaric Acid-gamma-lactone | 0.16 | 0.18 | 0.13 | 0.33 | 0.24 |
| p-Hydroxy Phenyl Acetic Acid | 0.16 | 0.17 | 0.13 | 0.58 | 0.01 |
| i-Erythritol | 0.19 | 0.17 | 0.15 | 0.26 | 0.05 |
| Beta-Hydroxy Butyric Acid | 0.20 | 0.19 | 0.15 | 0.91 | 0.16 |
| Sucrose | 0.16 | 0.17 | 0.13 | 0.65 | 0.22 |
| D-Threonine | 0.14 | 0.15 | 0.13 | 0.63 | 0.51 |
| D-Ribono-1,4-Lactone | 0.16 | 0.16 | 0.15 | 0.96 | 0.26 |
| Tyramine | 0.16 | 0.17 | 0.13 | 0.57 | 0.18 |
| Glycolic Acid | 0.15 | 0.17 | 0.13 | 0.16 | 0.10 |
| Chondroitin Sulfate C | 0.19 | 0.16 | 0.15 | 0.33 | 0.18 |
| L-Threonine | 0.16 | 0.18 | 0.13 | 0.44 | 0.08 |
| D-Saccharic Acid | 0.17 | 0.18 | 0.13 | 0.87 | 0.11 |
| D-Melibiose | 0.17 | 0.18 | 0.13 | 0.91 | 0.10 |
| 4-Hydroxy Benzoic Acid | 0.21 | 0.19 | 0.14 | 0.46 | 0.06 |
| D-Raffinose | 0.19 | 0.16 | 0.14 | 0.34 | 0.11 |
| Alpha-Keto-Glutaric Acid | 0.17 | 0.16 | 0.13 | 0.86 | 0.17 |
| Gelatin | 0.21 | 0.18 | 0.14 | 0.42 | 0.11 |
| L-Lysine | 0.18 | 0.17 | 0.14 | 0.73 | 0.26 |
| Stachyose | 0.19 | 0.18 | 0.14 | 0.65 | 0.19 |
| L-Serine | 0.17 | 0.17 | 0.13 | 0.96 | 0.01 |
| Mono Methyl Succinate | 0.16 | 0.17 | 0.13 | 0.39 | 0.01 |
| Laminarin | 0.17 | 0.18 | 0.14 | 0.92 | 0.15 |
| Glycyl-L-Aspartic Acid | 0.18 | 0.17 | 0.13 | 0.56 | 0.14 |
| D-Aspartic Acid | 0.15 | 0.16 | 0.13 | 0.61 | 0.17 |
| Lactulose | 0.17 | 0.21 | 0.13 | 0.42 | 0.15 |
| L-Glutamic Acid | 0.17 | 0.18 | 0.13 | 0.74 | 0.06 |
| Tricarballic Acid | 0.15 | 0.17 | 0.13 | 0.31 | 0.09 |
| Citraconic Acid | 0.18 | 0.18 | 0.14 | 0.95 | 0.27 |
| 1,2-Propanediol | 0.16 | 0.17 | 0.13 | 0.81 | 0.25 |
| L-Valine | 0.17 | 0.17 | 0.14 | 1.00 | 0.27 |
| Formic Acid | 0.15 | 0.17 | 0.13 | 0.35 | 0.10 |
| Phenylethylamine | 0.15 | 0.16 | 0.13 | 0.39 | 0.30 |
| N-Acetyl-L-Glutamic Acid | 0.18 | 0.17 | 0.14 | 0.87 | 0.22 |
| Fumaric Acid | 0.15 | 0.17 | 0.13 | 0.30 | 0.08 |
| Citric Acid | 0.14 | 0.16 | 0.13 | 0.29 | 0.19 |
| Succinamic Acid | 0.18 | 0.16 | 0.14 | 0.55 | 0.17 |
| N-Acetyl-D-Galactosamine | 0.16 | 0.16 | 0.14 | 0.96 | 0.38 |
| Glycyl-L-Glutamic Acid | 0.17 | 0.17 | 0.13 | 0.75 | 0.03 |
| L-Homoserine | 0.17 | 0.16 | 0.14 | 0.71 | 0.16 |
| Tween 20 | 0.15 | 0.14 | 0.12 | 0.91 | 0.44 |
| L-Glutamine | 0.15 | 0.16 | 0.12 | 0.18 | 0.07 |
| L-Isoleucine | 0.17 | 0.16 | 0.14 | 0.92 | 0.31 |
| myo-Inositol | 0.15 | 0.17 | 0.12 | 0.15 | 0.08 |
| L-Ornithine | 0.17 | 0.17 | 0.14 | 0.94 | 0.10 |
| 2,3-Butanediol | 0.19 | 0.17 | 0.14 | 0.58 | 0.12 |
| Glycolic Acid | 0.17 | 0.17 | 0.14 | 0.93 | 0.18 |
| D-Serine | 0.15 | 0.14 | 0.12 | 0.90 | 0.13 |
| Itaconic Acid | 0.16 | 0.14 | 0.14 | 0.47 | 0.36 |
| L-Asparagine | 0.16 | 0.17 | 0.12 | 0.79 | 0.12 |
| Hydroxy-L Proline | 0.18 | 0.17 | 0.14 | 0.56 | 0.10 |
| Quinic Acid | 0.18 | 0.17 | 0.13 | 0.65 | 0.03 |
| Melibionc Acid | 0.23 | 0.20 | 0.13 | 0.82 | 0.32 |
| gamma-Amino Butyric Acid | 0.17 | 0.17 | 0.13 | 0.97 | 0.24 |
| D,L-Malic Acid | 0.16 | 0.17 | 0.12 | 0.58 | 0.06 |
| D-Glucosaminic Acid | 0.15 | 0.17 | 0.12 | 0.21 | 0.10 |
| L-Malic Acid | 0.15 | 0.15 | 0.12 | 0.39 | 0.00 |
| L-Arginine | 0.17 | 0.16 | 0.13 | 0.48 | 0.11 |
| D,L-Carnitine | 0.17 | 0.17 | 0.13 | 0.88 | 0.10 |
| Acetamide | 0.18 | 0.17 | 0.13 | 0.71 | 0.10 |
| D-Tartaric Acid | 0.17 | 0.17 | 0.13 | 0.94 | 0.14 |
| Citramalic Acid | 0.17 | 0.17 | 0.13 | 0.95 | 0.17 |
| L-Phenylalanine | 0.17 | 0.16 | 0.13 | 0.85 | 0.11 |
| D-Lactic Acid Methyl Ester | 0.16 | 0.16 | 0.13 | 0.97 | 0.07 |
| L-Tartaric Acid | 0.17 | 0.17 | 0.13 | 0.88 | 0.02 |
| Malonic Acid | 0.17 | 0.16 | 0.12 | 0.52 | 0.10 |

